# Lesions of anterior cingulate cortex disrupt an electrophysiological signature of reward processing in humans

**DOI:** 10.1101/2024.12.04.626789

**Authors:** Joyce Oerlemans, Ricardo J. Alejandro, Dimitri Hemelsoet, Eva Genbrugge, Katie Bouche, Luc Defreyne, Veerle De Herdt, Clay B. Holroyd

**Affiliations:** 4BRAIN, Department of Head and Skin, Ghent University, 9000 Ghent, Belgium; Department of Experimental Psychology, Ghent University, 9000 Ghent, Belgium; Department of Neurology, Ghent University Hospital, 9000, Belgium; Department of Radiology, Ghent University Hospital, 9000, Belgium; Department of Physical and Rehabilitation Medicine, Ghent University Hospital, 9000, Belgium; Department of Vascular and Interventional Radiology, Ghent University Hospital, 9000, Ghent

**Keywords:** anterior cingulate cortex, reward positivity, voxel-based lesion-symptom mapping

## Abstract

The reward positivity (RewP) is an event-related brain potential (ERP) component associated with feedback and reward processing. Although the component is said to be generated in anterior cingulate cortex (ACC), this inference is disputed because of the inverse problem. Recently, by conducting a current source density analysis of intracranial electroencephalogram (EEG) data recorded from a large cohort of epilepsy patients, we provided direct evidence that the RewP is produced by a circumscribed region in caudal ACC corresponding to Brodmann areas 24c’ and 32’. In the present study we confirm that this brain area is the source of the RewP by examining the effects of damage to frontal cortex on RewP amplitude. We recorded scalp EEG from 68 stroke patients with damage to frontal cortex while they engaged in a trial-and-error guessing task used to elicit a canonical RewP. Application of non-parametric voxel-based lesion-symptom mapping to the lesion data revealed that damage to Brodmann areas 24c’ and 32’ attenuated RewP amplitude, whereas damage to other parts of frontal cortex did not affect it. These results provide causal evidence that the caudal ACC generates the RewP and underscore the contribution of this brain region toward motivating extended behaviours.

**Significance statement:** The reward positivity (RewP) is an event-related brain potential component that reflects reward processing in the brain. Although multiple studies have suggested that anterior cingulate cortex (ACC) is the neural generator of the RewP, this assertion is disputed mainly because of the inverse problem. In this study, we present direct evidence that the RewP is generated in caudal ACC by performing voxel-based lesion-symptom mapping (VLSM) in a cohort of 68 stroke patients with lesions in the frontal lobe. We found that a circumscribed region in right caudal ACC was significantly associated with a reduced RewP, providing the first causal evidence that this region generates the RewP in humans. These fundings elucidate the central role ACC plays in motivating extended behaviours.

## Introduction

The anterior cingulate cortex (ACC) is a broad cortical region located bilaterally in the medial portion of the prefrontal cortex ^1^. Although it is generally accepted that ACC plays a central role in cognitive control, its exact function remains in dispute ^2,3^. Converging evidence indicates that ACC regulates effort expenditure according to the predicted reward value of the task at hand ^4,5^. In turn, the reward-prediction function of ACC has been inferred partly by the discovery of a component of the human event-related brain potential (ERP) called the reward positivity (RewP). This component, which is proposed to be generated by ACC ^6^, is elicited over frontal-central areas of the scalp about 300 milliseconds (ms) following presentation of feedback stimuli in trial-and-error learning and guessing tasks ^7,8^. Moreover, the RewP is larger to unexpected rewards than to unexpected non-rewards ^7,9^, consistent with the proposal that it encodes a reward prediction error important for reinforcement learning and value-driven decision making ^10,11^. These observations have motivated abundant research indicating that the RewP is sensitive to a neural mechanism for reward processing ^12^, and by extension, that ACC serves to motivate the execution of complex behaviours ^13,14^.

Nevertheless, the exact neural origins of the component have been debated for years ^15^. Initial source-localization efforts based on the scalp electroencephalogram (EEG) suggested a generator in ACC ^6,16– 20^, but other studies have found sources for the RewP in the posterior cingulate cortex ^21–23^ and the basal ganglia ^24^. Still other studies using fMRI ^25^, combined EEG/fMRI ^26–28^, and intracranial recordings in rodents ^29^, non-human primates ^30^, and humans ^31^ have also implicated the ACC as a source of the RewP, but these findings are limited by their respective methodologies (including low temporal resolution of the fMRI blood-oxygen-level-dependent signal ^32^, imperfect homologies between animal and human brains, and non-standard methods for assessing RewP amplitude). In general, identifying the neural sources of any EEG signal from only the scalp EEG is unsolvable because the problem is mathematically ill-posed (relating to the so-called inverse problem, which asserts that an infinite number of neural source configurations can produce the same EEG signal recorded at a specific scalp electrode) ^33^.

For this reason, we recently addressed this question directly by recording intracranial EEG-data across a large cohort of epilepsy patients ^34^. After pooling the data across subjects, we compared the current-source densities of neural activity across multiple brain regions. This analysis revealed that a relatively circumscribed area within caudal ACC spanning the cingulate and paracingulate sulci, including Brodmann areas 24c’ and 32’ (also called anterior midcingulate cortex ^1,35^, Supplementary Figure 1), best accounted for the time course of the RewP observed at the scalp. Further, this region coincided with an area of peak feedback-related activity observed in a previous fMRI experiment ^25^, and is where more than two decades ago was predicted that the RewP would be generated ^10^.

Nevertheless, although the CSD activity observed in caudal ACC was strongly correlated with the RewP, the question about whether this brain region causally produces the RewP remains open. Here we addressed this question by investigating the consequences of damage to frontal cortex on RewP amplitude. We hypothesized that if caudal ACC produces the RewP, then damage to this brain area would attenuate RewP amplitude. To test this hypothesis, we recruited 72 patients with stroke damage involving any region of the frontal lobe, including lesions in caudal and rostral ACC. The participants engaged in a task that generates a canonical RewP ^9^ while their scalp EEG was recorded. Non-parametric voxel-based lesion-symptom mapping (VLSM) was applied to identify frontal regions where brain damage was associated with reduced RewP amplitude ^36^. Confirming our prediction, we found that damage to the caudal ACC, where we previously observed the RewP to be produced, uniquely resulted in a smaller RewP.

## Materials and methods

### Ethics statement

The Ethics Committee of Ghent University Hospital reviewed and approved this study (BC-08764). All patients provided written informed consent before participating in this research. This study was conducted in accord with the Declaration of Helsinki.

### Participants

Seventy-two stroke patients with focal lesions in the frontal lobe were enrolled prospectively and recruited at three departments of Ghent University Hospital: the Department of Neurology, the Department of Vascular and Interventional Radiology, and the Rehabilitation Center. Inclusion criteria were 1) focal lesions in the frontal lobe due to ischemic stroke, haemorrhagic stroke or subarachnoid haemorrhage from arteria cerebri anterior (ACA) aneurysms, and 2) recent neuroimaging evidence (either CT or MRI, maximal 2 years old) confirming damage to frontal cortex. We recruited patients both during the acute and chronic phase after stroke onset to maximize recruitment rate. Exclusion criteria were 1) a history of neurodegenerative diseases or psychiatric disorders, 2) history of lesions in the frontal lobe, and 3) active substance abuse. Patients with lesions that extended from the frontal lobe into other parts of the brain were not excluded from the study.

### Task procedure

Participants first performed a hierarchical action sequence task, the results of which will be presented in Study III (Chapter VII). Then, following a break during which EEG electrodes were applied, the participants performed a virtual T-maze task (vTMT), a task that is known to elicit a robust RewP ^9^. Participants were required to navigate a simulated maze by choosing a left or right response at the start of each trial, as indicated by the appearance of a green arrow in the center of the stem image at the maze intersect. At the time of their response, the image of the selected alley appeared for 500 ms, followed by the feedback of a fruit (more specifically an apple or an orange). The alley and the fruit remained together on the screen for 1000 ms. Participants were instructed that presentation of one type of fruit indicated that they chose the correct direction (reward/positive feedback), whereas the presentation of the other type of fruit indicated they chose the incorrect direction (no reward/negative feedback). The designation of positive or negative feedback to a specific type of fruit was counterbalanced across participants, and the type of feedback was selected at random (50% probability for each feedback type), but the participants were not informed of this. The experiment consisted of four blocks of 50 trials each, which resulted in a total of 200 trials per participant. For task details see Supplementary Figure 4 and Oerlemans et al. ^34^.

### ERP analysis

Scalp EEG was recorded using a Micromed 32 channel Brainquick mobile EEG system (Micromed, Italy) with a sampling rate of 512 Hz and an online high-pass filter of 0.15 Hz. 23 electrodes were placed according to the 10-20 system and 4 electrodes were placed near the eyes (2 horizontal, and 2 vertical). The right mastoid was used as the online reference electrode. All data were processed and visualized using BrainVision analyzer 2.2 (Brain Products GmbH). The digitized signals were filtered, segmented, corrected for artifacts, re-referenced and averaged according to the standard principles (see SI for details). The RewP was identified using a difference wave approach ^38^ by subtracting the average waveform of reward from the average waveform of no reward for every electrode and participant, resulting in an average difference wave (i.e., the RewP) for every electrode and participant ^37,52^.

Typically, the RewP reaches maximum amplitude at channel FCz ^37^. Because our recording system did not have an electrode available at that location, we constructed a virtual channel FCz by averaging the difference waves associated with channels Fz and Cz. RewP amplitude was assessed within a time window of 250 to 400 ms, which includes when the RewP usually occurs, but was wide enough to accommodate relatively large across-subject variability in RewP timing observed in this population ^11^. A wide window is of particular importance given that latency variability tends to be inversely related to ERP component amplitude, and thus might be expected to increase when the neural generator is damaged ^53^. Moreover, the average age of the participants in the stroke study population was around 60 years old, and RewP amplitude is known to decrease with age ^54^. RewP amplitude was measured for each subject in two steps. First, we identified the maximum negative amplitude of the difference wave (because the RewP is expressed as a negative-going deflection) at the virtual channel FCz within the pre-defined time window. If no negative deflection occurred within this time window (i.e., the difference wave was positive-going) or if the scalp distribution was posterior (as was the case for a single participant; see Supplementary Figure 5), then RewP amplitude was defined as 0 μV (in other words, this participant did not show a RewP). Second, for the subjects that showed a negative deflection within the time window, we determined the onset and offset of the RewP. RewP onset was taken as the time point of the first polarity reversal proceeding the RewP. If no reversal occurred within the pre-defined time window, then the lower bound was set at 250 ms (i.e., the edge of the window). An analogous procedure was conducted for the polarity reversal following the RewP to determine RewP offset. The voltages between RewP onset and RewP offset were then summed and divided by 78 (i.e., the number of samples within 250-400 ms) to determine RewP amplitude for each subject.

### Pre-processing of neuroanatomical data

A structural magnetic resonance imaging (MRI) scan for every patient was used to delineate lesion location and extent for each MRI slice. Lesion demarcation was based on either the DWI images when the MRI was performed less than 48 hours after stroke onset, or on the FLAIR images when MRI was performed more than 48 hours after stroke onset. The areas were manually traced by the first author using MRIcron to draw the regions of interest ^55^. Lesions that were difficult to identify in terms of their location and/or extension were discussed with an experienced neuroradiologist at Ghent University Hospital, who was blinded to the purpose of the study. The patient’s MRI scan and the corresponding lesion maps were co-registered to the anatomical T1-weighted sequence, and were subsequently spatially normalized to the Montreal Neurological Institute (MNI) standard space using SPM12 ^56^. Finally, an overlap map of the normalized lesion masks was created to assess lesion distribution and voxel-wise statistical power.

### Voxel-based lesion-symptom mapping

VLSM was applied to determine lesion locations that were significantly associated with smaller RewP amplitude ^36^. VLSM analyses were performed in the NiiStat toolbox for Matlab (Matlab 2021a, http://www.nitrc.org/projects/niistat). All analyses were corrected for age and lesion size using a general linear model (GLM) approach. Only voxels with a minimal overlap of 5% of the study population were included (n = 68, minimal overlap of 4 patients). For convenience, we defined RewP amplitude such that positive z-scores corresponded to reduced RewP amplitude. VLSM analyses were run separately for left and right cerebral hemisphere. To test our hypothesis that the RewP is produced in a small region of ACC associated with RewP activity and feedback processing ^25,34^, we examined 5 ROIs spanning this area as described by the AICHA atlas available in NiiStat ^39^. These included 5 regions that spanned the predicted ROI: cingulate sulci 1 and 2 (S_cingulate 1 and 2), midcingulate gyrus 1 (G_cingulum_mid 1), superior medial frontal gyrus 3 (G_frontal_sup_medial 3), and supplementary motor area 1 (G_sup_motor_area 1) (Figure 4). To explore whether damage to other regions in the frontal lobe might also disrupt the RewP – even though this was not our hypothesis – we conducted two follow-up analyses based on the AAL atlas ^40^ of the frontal medial wall specifically and the entire frontal lobe generally.

## Results

We recruited 72 patients with lesions in the frontal lobe, 68 of whom (94.4%) successfully completed the virtual T-maze task and were included for further data analysis (see Supplementary Figure 2 for flowchart). The mean age was 60.4 years old (±14, the youngest patient was 18, the oldest 85), and 47.1% (n=32) were female. Of those 68 patients, 17 patients (25%) had lesions in the ACC, 8 of which (47.1%) were located in the rostral ACC area, and the remaining 9 (52.9%) were located in the caudal ACC area (see supplementary Table S1 for demographic details). The overlay map of the stroke lesions of all patients indicated that the lesions were distributed mainly in the frontal lobe, but also in the parietal, temporal, and occipital lobes (Figure 1).

**Figure 1.**
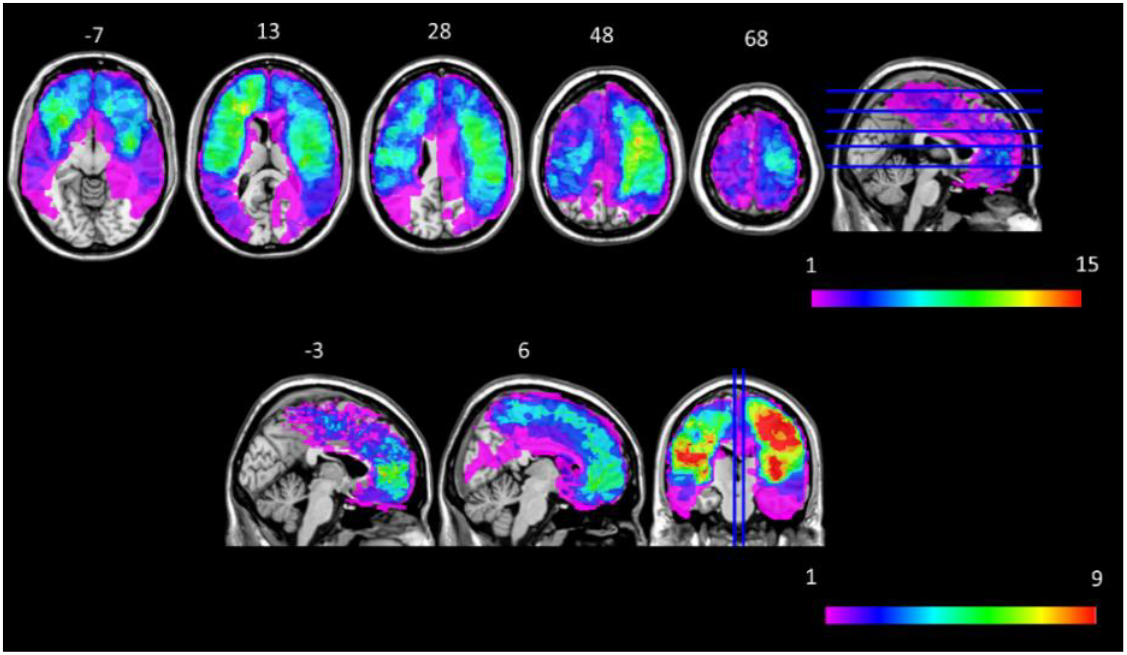
Map of individual stroke lesions of all 68 patients overlaid on the ICBM 152 template in Montreal Neurological Institute (MNI) space. Top: axial view (except for the last image which shows a sagittal view of the presented axial slices). The colour scale indicates the number of patients having a lesion at each voxel (ranging from 1 to 15 patients, which was the highest overlap). The MNI coordinates of each transverse section (z-axis) are given above each image. Bottom: sagittal view of the left and right frontal medial wall (except for the last image which shows a coronal view of the presented sagittal slices). The colour scale indicates the number of patients having a lesion at each voxel (ranging from 1 to 9 patients, which was the highest overlap in the medial frontal wall). The MNI coordinates of each sagittal section (x-axis) are given above each image.

Figure 2A displays the grand average (GA) RewP for the entire study population (n=68). Note that the RewP is typically calculated by measuring the difference between ERP responses elicited by positive vs. negative feedback ^6,37^. This approach isolates neural processes related to feedback valence by removing sources of variance that are common to both conditions (i.e., the difference wave approach) ^37,38^. The RewP difference wave shows a peak at 318 ms after feedback stimulus onset (±44.5, 95% confidence interval [315, 335]), with a peak amplitude of -1.1 µV (±0.31, 95% confidence interval [- 0.82, -0.68]), and exhibited a frontocentral scalp distribution that was slightly lateralized to the right (Fig 2A, inset).

**Figure 2.**
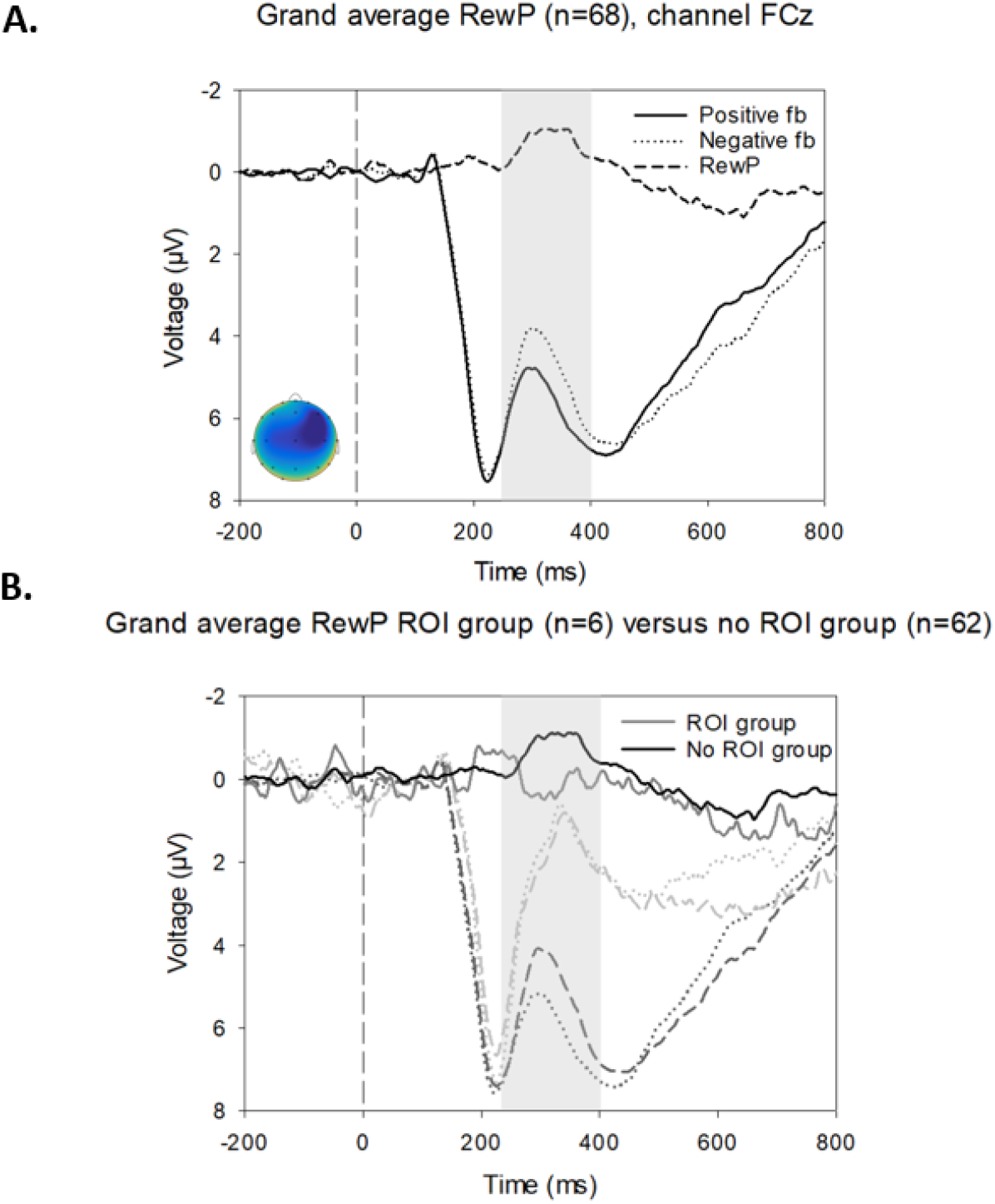
The reward positivity. A. The grand average event-related potentials (ERPs) to positive and negative feedback stimuli and reward positivity difference wave of all patients who successfully completed the vTMT (n=68), recorded at virtual channel FCz. Solid line = average ERP after positive feedback, dotted line = average ERP after negative feedback, striped line = difference wave (negative feedback – positive feedback). The RewP is identified as the negative-going activity in the difference wave (plotted upwards according to convention). The peak of the RewP occurred at 318.6 ms after feedback presentation (vertical dashed line). The scalp distribution of the grand average (inset) shows a frontocentral scalp distribution (blue = maximal negative value -1.18 µV, yellow = minimal negative value -0.35 µV). B. Grand average of the difference waveform (negative feedback – positive feedback) for the patients with lesions in the ROI (‘ROI’ group, n=6) versus patients with lesions outside the ROI (‘no ROI’ group, n=62) for data recorded at channel FCz. Dashed lines = average ERPs after negative feedback, dotted lines = average ERPs after positive feedback, solid lines = difference waves. ‘ROI group’ refers to the group of patients with lesions in the right caudal ACC ROI; ‘no ROI’ group refers to the group of patients with lesions outside of the right caudal ACC ROI. The RewP time window (250 – 400 ms) is highlighted in grey.

The relationship between lesion location and RewP amplitude was analysed using VLSM (statistically corrected for age and lesion size). Figure 3 shows the uncorrected thresholds maps reported as z-scores; positive scores correspond to reduced RewP amplitude. Visual inspection of the figure suggests that damage of right ACC is associated with smaller RewP amplitude.

**Figure 3.**
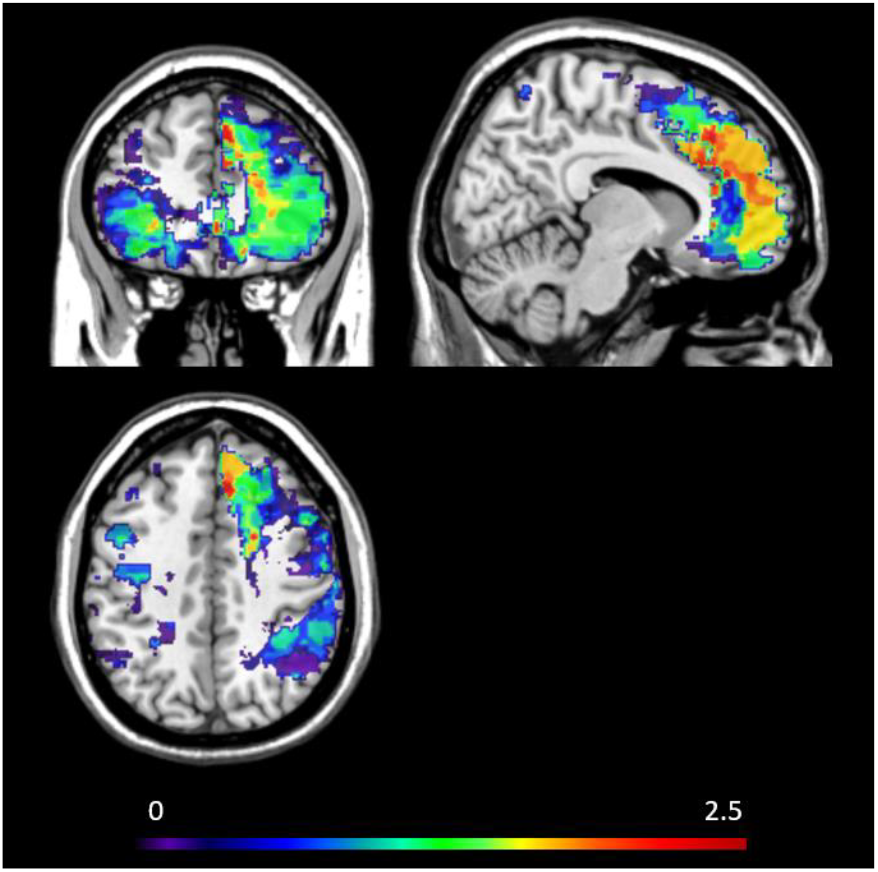
Uncorrected threshold map displaying the results of voxel-based lesion-symptom mapping with reward positivity amplitude as a dependent variable, corrected for age and lesion volume. The highest z-score in the data was associated with right caudal ACC (z=2.4, red area).

To test our hypothesis that the RewP is generated in the region of caudal ACC that we identified in our previous study ^34^, we applied VLSM to the lesion data within five regions of the AICHA atlas ^39^ that impinged on this region of interest (ROI) (Figure 4). Analyses for the left and right cerebral hemispheres were performed independently. Two regions in the right hemisphere were statistically significant: the right frontal superior medial cortex (z=1.96) and the right cingulate cortex 1 (z=1.85) (Figure 4). For the purpose of illustration, the grand averages for the group of patients who contributed to the right caudal ACC ROI 25,34 (n=6) and for the group of patients who had lesions outside of the ROI (n=62) are presented in Figure 2B. The group of patients with lesions outside the ROI produced a normal RewP having an amplitude of -0.81 µV (Figure 2B, No ROI group). By contrast, the group of patients with lesions impinging on the ROI produced a RewP that was much smaller with an amplitude of -0.05 µV (Figure 2B, ROI group). These results confirm our prediction that damage to caudal ACC would attenuate RewP amplitude.

Additionally, we explored whether damage to brain areas outside of the ROI would also attenuate RewP amplitude. We conducted two analyses using the AAL atlas ^40^, which provides more course-grain ROIs than does the AICHA atlas. The first analysis, which examined 4 ROIs encompassing the entire frontal medial wall (including the ROI tested in the main analysis), revealed that damage to the right super medial frontal cortex was significantly associated with smaller RewP amplitude (see Supplementary Figure 3). This result is consistent with our overall conclusion that the RewP is produced within caudal ACC. The second analysis, which examined 10 ROIs that include most of the frontal lobe, did not yield statistically significant results (Supplementary Figure 3).

**Figure 4.**
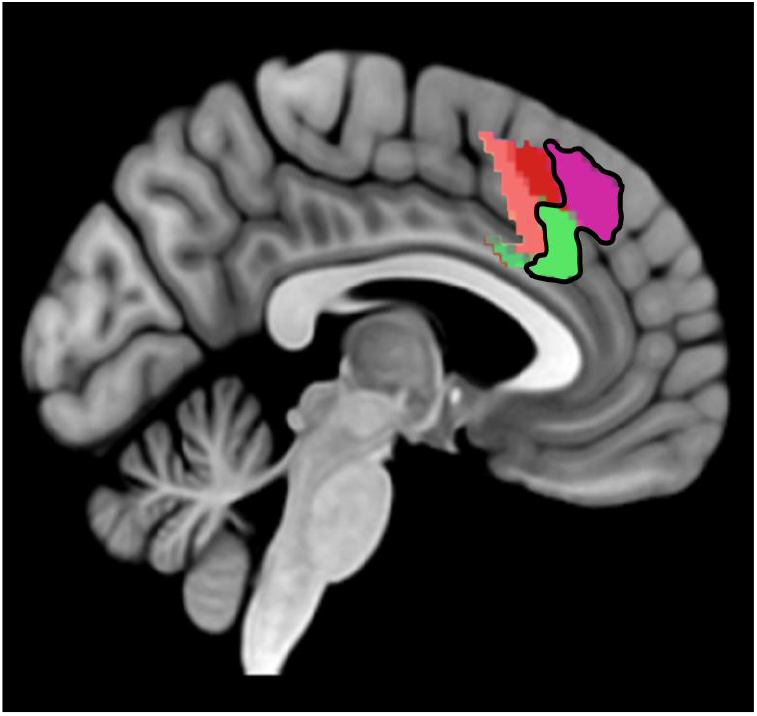
Five regions of the AICHA atlas ^39^ selected for our VLSM analysis of caudal ACC overlaid on the MNI brain template. The 5 ROIs that were included in the VLSM analysis were: S cingulate 1 (green), S cingulate 2 (salmon pink), G supplementary motor area (red), G frontal superior medial cortex (purple), and G cingulum mid (dark green, on the posterior side of S cingulate 1). The 2 ROIs that survived the permutation analysis are encircled in black: right G frontal superior medial cortex (z=1.96, purple) and right S cingulate 1 (z=1.85, green). Damage to these two ROIs was significantly associated with a reduced RewP amplitude.

## Discussion

We found that damage to right caudal ACC was associated with reduced RewP amplitude, which is an ERP component that occurs about 250 to 350 ms after presentation of reward-related feedback stimuli ^6^ indicating whether ongoing events are better or worse than expected ^10,41^. Although the RewP has historically been attributed to neural activity occurring in ACC, this claim is controversial due to the inverse problem ^33^. Here we conducted a loss-of-function study involving 68 patients with damage to the frontal lobes as they engaged in a guessing task with feedback while their EEG was recorded ^36,42^. VLSM analysis revealed that damage to a region spanning right caudal cingulate and paracingulate gyri including Brodmann areas 24c’ and 32’ ^34^ was associated with reduced RewP amplitude. This region corresponds to a brain area that we previously identified as the RewP generator in a current source density analysis of intracranial EEG recordings pooled across multiple human epilepsy patients. Our results converge with the previous study and provide the first causal evidence that caudal ACC is the neural generator of the RewP in humans.

These findings also raise several questions. In particular, the effect of cortical damage on RewP amplitude was limited to the right cerebral hemisphere, whereas the results of our previous study were lateralized to the left cerebral hemisphere ^34^. Multiple explanations could account for this discrepancy, but it seems likely that the left caudal ACC lacked sufficient statistical power to exhibit an effect were it to exist. Indeed, the overlap density is higher in the right caudal ACC ROI (4 patients) than in the left caudal ACC ROI (3 patients), which was below the threshold for inclusion. This shortcoming exemplifies the major challenges of loss-of-function studies, namely, obtaining large sample sizes that yield sufficient statistical power ^36,42^. This challenge is especially pronounced for studies that focus on ACC specifically, as this brain region is vascularized by the anterior cerebral artery (ACA) ^43^ and isolated lesions in the ACA territory are very rare ^44^. Although subarachnoid haemorrhages due to rupture of an ACA aneurysm are more prevalent ^45^, these are not always accompanied by actual lesions of the brain tissue. Additionally, the morphology of the cingulate and paracingulate cortex exhibits significant interindividual variability ^46–48^, which has been demonstrated to affect reward feedback processing in these regions ^25^.

To maximize the number of participants, we not only recruited patients from different departments in the hospital (i.e., the Department of Neurology, the Department of Vascular and Interventional Radiology, and the Rehabilitation Center), but we also recruited patients irrespective of the phase of the stroke. There has been an ongoing debate about the optimal timing for recruiting stroke patients to study cognition. On one hand, during the acute phase (i.e., during the first week after the stroke) performance on cognitive tasks can be influenced by possible generic effects of the stroke such as headache or tiredness, and the lesion might still be evolving ^49^. On the other hand, during the chronic phase (usually starting from 6 months) some re-organization of the neural pathways might already have occurred, compensating for the focal effects of brain damage on cognition ^42^. Because of concerns about obtaining a sufficiently large sample size for this study, the patients were asked to participate irrespective of the phase of occurrence. It is possible that RewP amplitude differed between the acute and chronic phases in ways that could have impacted the results, which is a weakness of the study design.

Because the VLSM approach tests for the effects of lesion damage on the dependent variable independently for each voxel, correction for multiple comparisons can severely limit statistical power. A widely-employed mitigation strategy is to group collections of voxels into ROIs, which minimizes the number of multiple comparisons ^42,50^. The ROI approach is especially recommended for studies with a clear a priori hypothesis, as was the case for our study. In particular, our ROI was based on a previous study in which we demonstrated that the time course of the RewP measured at the scalp was strongly correlated with the dynamics of current-source-density activity in caudal ACC ^34^, providing evidence of source generation ^51^. This particular study also utilized an ROI from a previous fMRI experiment that demonstrated that feedback activity was associated with individual differences in the anatomy of the cingulate and paracingulate cortices within Brodmann areas 24c’ and 32’ ^25^. These previous findings justify the use of this brain area as the ROI for the present study.

Further, to explore whether damage to other areas of the frontal lobe also impact the RewP, we conducted two additional ROI analyses using the AAL atlas ^40^, which is composed of larger ROIs. A VLSM analysis of 4 large ROIs across the frontal medial wall revealed a significant association between the right medial superior frontal cortex and reduced RewP amplitude, consistent with our first ROI analysis. A second VLSM analysis including 10 ROIs distributed across the entire frontal lobe did not reveal any significant associations. Notably, cortical damage was distributed widely throughout frontal cortex (Figure 1), so the lack of an association between RewP amplitude and lesions outside of medial frontal cortex is unlikely to be due to statistical biases in the raw data. As evidence of this, in 37 patients from the same study who also completed a behavioural hierarchical sequence task, VLSM indicated that lesions of right DLPFC impaired performance of that task (findings presented in Study III). These results are indicative of sufficient statistical power in our study population to detect neural and behavioural effects elsewhere in frontal cortex, even in a subsample consisting of about half of the participants in our study.

In summary, the results of our study confirm that the RewP is produced in a region of caudal ACC centered around the cingulate sulcus and extending to the paracingulate sulcus (specifically, areas 24c’ and 32’) ^1,35^. Based on considerations of functional neuroanatomy of ACC and the midbrain dopamine system, which conveys reward prediction error signals to frontal cortex, we predicted more than two decades ago that the RewP is produced in the ventral bank of the cingulate sulcus, namely in Brodmann area 24c’ ^10^. In recent fMRI work we have also found that this area is sensitive to reward-related modulation of distributed representations of task progress in the production of hierarchically-organized action sequences. Our present results support these findings by highlighting a hotspot of reward processing in ACC that motivates the execution of extended behaviours ^13^.

## Acknowledgments

The authors would like to thank all the participating patients who made this study possible. We would also like to acknowledge Travis Baker for providing the virtual T-maze task code. This work is part of a project that has received funding from the European Research Council (ERC) under the EU’s Horizon 2020 Research and Innovation Program (grant agreement no. 787307).

