## Supplementary Materials for "Lesions of anterior cingulate cortex disrupt an electrophysiological signature of reward processing in humans"

### Supporting Information Text: pre-processing of scalp EEG.

Scalp EEG was recorded on a Micromed 32 channel Branquick mobile EEG system (Micromed, Italy) with a sampling rate of 512 Hz and an online high-pass filter of 0.15 Hz, using 23 electrode sites and 4 eye electrodes (2 horizontal, 2 vertical). The right mastoid electrode was used as the online reference electrode. All data were processed and visualized using BrainVision Analyzer 2.2 (Brain Products GmbH). The digitized signals were filtered with a zero-phase shift Butterworth filter using a passband of 0.1-20 Hz. A 1000 ms epoch of data extending from 200 ms prior to 800 ms following the onset of each feedback stimulus was extracted from the continuous data file for analysis. Ocular artifacts were corrected using the eye movement correction algorithm described by Gratton et al.[57]. The EEG data were re-referenced to the average of the left and right mastoids and were baseline corrected by subtracting from each sample the mean voltage associated with that electrode during the 200 ms interval preceding stimulus onset. Other artifacts were removed using a ± 100 microvolt (μV) level threshold and a 35 μV step threshold as rejection criteria, using a semi-automatic approach (meaning that the software identified trials to be rejected, but the researcher also inspected each trial visually for the final decision about its inclusion or exclusion). ERPs were created for each electrode and participant by averaging the single-trial EEG separately according to the feedback condition (reward, no reward). The RewP was identified using a standard difference wave approach by subtracting the average waveform of reward from the average waveform of no reward for every electrode and participant, resulting in an average difference wave (i.e., the RewP) for every electrode and participant [38].

### Supplementary Figure 1: Caudal ACC ROI.

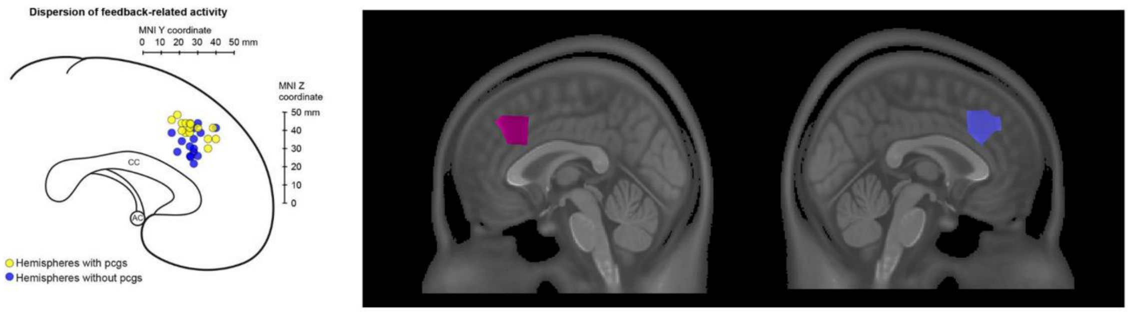


**Supplementary Fig 1.** Caudal ACC ROI in the Amiez et al. study [25] (left panel) and the Oerlemans et al. study [34] (right panel). The pink ROI refers to an area that impinges on the caudal cingulate and paracingulate sulcus on the left cerebral hemisphere; the blue ROI refers to the equivalent ROI on the right cerebral hemisphere.

**Supplementary Figure 2: Flowchart of patient selection.**

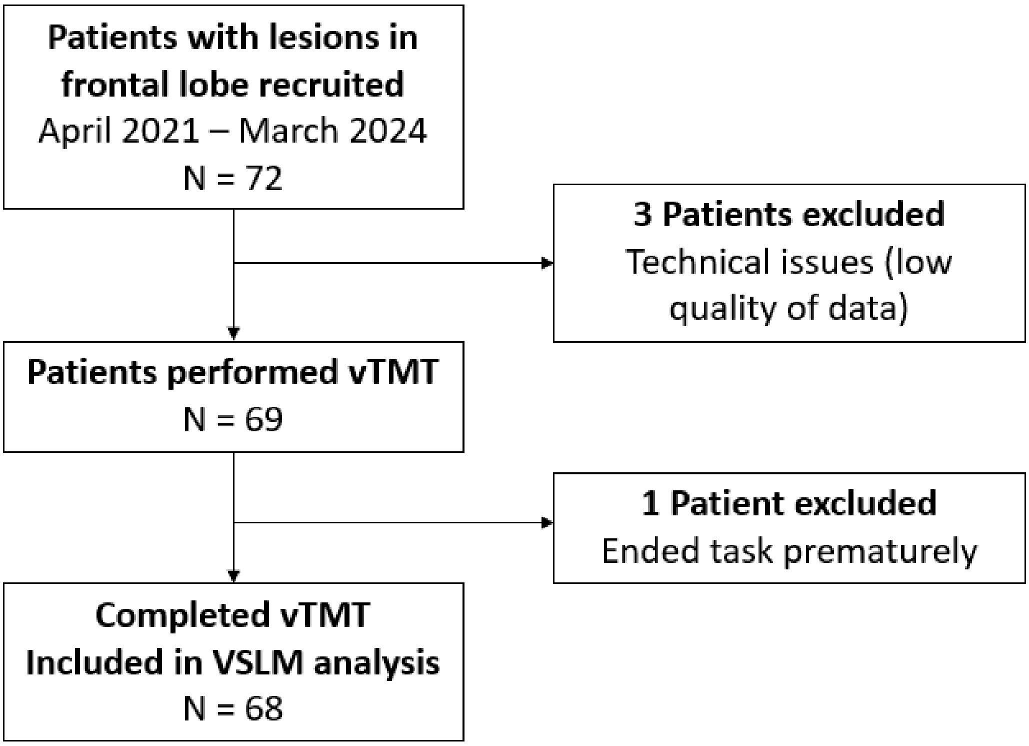


**Supplementary Figure 3: Regions of interest for VLMS analysis.**

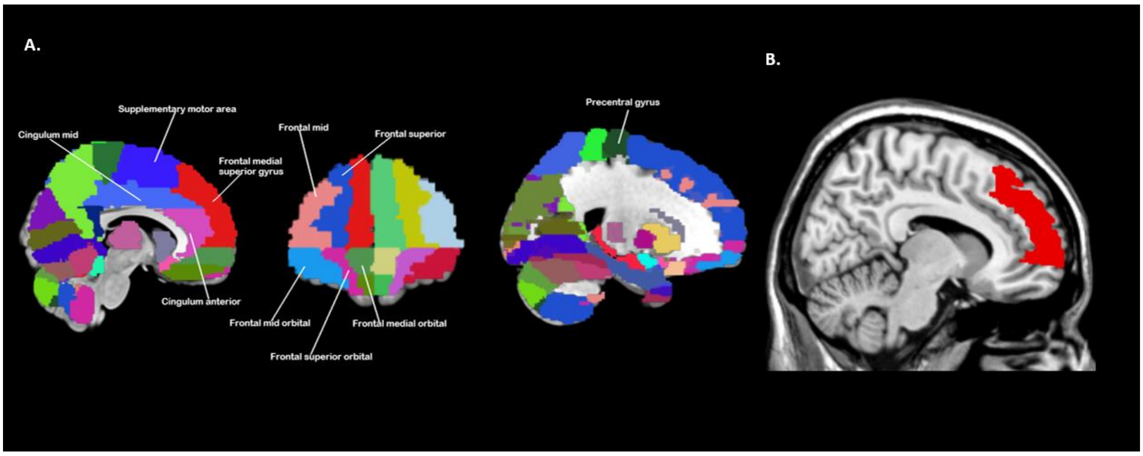


**Supplementary Fig. 3**. ROIs included in VLSM analysis for entire frontal medial wall and the entire frontal lobe based on the AAL atlas [40]. A. The ROIs that constituted the medial frontal wall were: supplementary motor area, frontal superior medial gyrus, cingulum mid and cingulum anterior. We added six more ROIs to cover the entire frontal lobe, namely frontal mid orbital, frontal superior orbital, frontal medial orbital, frontal mid, frontal superior and precentral gyrus [40]. B. VLSM analysis for the entire frontal medial wall revealed a statistically significant association between damage to the right frontal medial superior cortex (red) and a reduced RewP amplitude.

### Supplementary Figure 4: Overview of the virtual T-maze task.

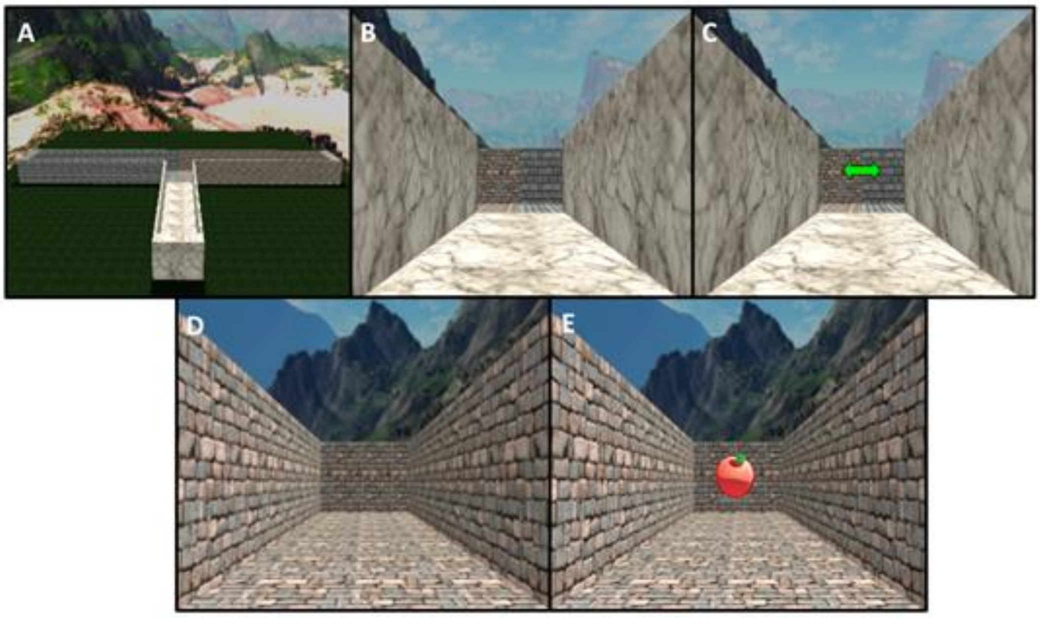


**Supplementary Fig 4.** A. View of the T-maze from above. B. Starting point of new trials (appears for 1000 ms). C. The double green arrow appears, indicating to the participant to choose a direction (left or right). The arrow remains visible until the button press. D. View of the selected alley (in this case left) (remains on screen for 500 ms). E. Image of the feedback stimulus (apple or orange), indicating that the participant chose the correct or incorrect direction (shown for 1000 ms).

### Supplementary Figure 5: Excluded participant.

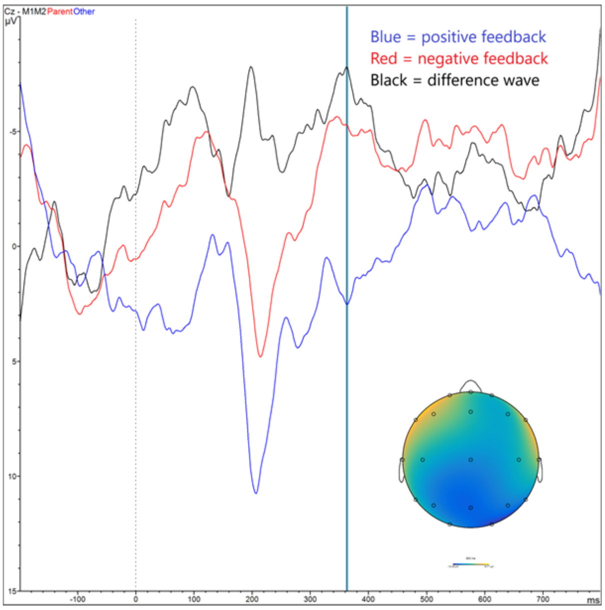


**Supplementary Fig 5.** Participant with posterior scalp distribution, recorded at channel Cz. Scalp distribution of the difference wave at 363 ms after reward feedback (i.e., peak within the RewP time window) is posterior. Range of the scalp distribution presented: -10.33 µV (dark blue); -0.71 µV (yellow). Only 97 of the 200 trials were withheld for this patient, due to excessive noise. The RewP value was set at 0 µV.

**Supplementary Table 1: Additional demographic information about the study population.**

| **Patient characteristics** | |
|---|---|
| ***Age*** | Mean 60.38<br>Median 62<br>SD 13.978<br>Min 18 – Max 85 |
| ***Gender*** | 47.1 % female, 52.9% male |
| ***Stroke information*** | 69.1% ischemic, 30.9% haemorrhagic |
| ***Lesion location*** | 25% ACC lesion (n=17)<br>11.8% rostral ACC (n=8)<br>13.2% caudal ACC (n=9) |
| ***Side of ACC lesion*** | Left (n=8)<br>Right (n=7)<br>Bilateral (n=2) |
| ***RewP AUC*** | Mean 1.17681<br>Median 0.8227<br>SD 1.1940228<br>Min 0 – max 4.4737 |
| ***Patients who contributed to caudal ACC ROI*** | Left hemisphere n=9<br>Right hemisphere n=8 |
